# Precise, high-throughput production of multicellular spheroids with a bespoke 3D bioprinter

**DOI:** 10.1101/2020.04.06.028548

**Authors:** Robert H. Utama, Lakmali Atapattu, Aidan P. O’Mahony, Christopher M. Fife, Jongho Baek, Théophile Allard, Kieran J. O’Mahony, Julio Ribeiro, Katharina Gaus, Maria Kavallaris, J. Justin Gooding

**Author notes:** Equal contributions: R.H.U; L.A. Maria Kavallaris and J. Justin Gooding, (M.K.); (J. J.G.).

## Abstract

3D *in vitro* cancer models are important therapeutic and biological discovery tools, yet formation of multicellular spheroids in a throughput and highly controlled manner to achieve robust and statistically relevant data, remains challenging. Here, we developed an enabling technology consisting of a bespoke drop-on-demand 3D bioprinter capable of high-throughput printing of 96-well plates of spheroids. 3D-multicellular spheroids are embedded inside a tissue-like matrix with precise control over size and cell number. Application of 3D bioprinting for high-throughput drug screening was demonstrated with doxorubicin. Measurements showed that IC_50_ values were sensitive to spheroid size, embedding and how spheroids conform to the embedding, revealing parameters shaping biological responses in these models. Our study demonstrates the potential of 3D bioprinting as a robust high-throughput platform to screen biological and therapeutic parameters.

**Significance Statement:** *In vitro* 3D cell cultures serve as more realistic models, compared to 2D cell culture, for understanding diverse biology and for drug discovery. Preparing 3D cell cultures with defined parameters is challenging, with significant failure rates when embedding 3D multicellular spheroids into extracellular mimics. Here, we report a new 3D bioprinter we developed in conjunction with bioinks to allow 3D-multicellular spheroids to be produced in a high-throughput manner. High-throughput production of embedded multicellular spheroids allowed entire drug-dose responses to be performed in 96-well plate format with statistically relevant numbers of data points. We have deconvoluted important parameters in drug responses including the impact of spheroid size and embedding in an extracellular matrix mimic on IC_50_ values.

## Introduction

3D cell cultures have a number of advantages over 2D cell cultures and should be the workhorse of modern cell biology (1). For example, in cancer cell biology, a number of cellular properties, such as cell proliferation, differentiation and the response to drugs are fundamentally different for cells in 2D and 3D environments (2, 3). Further, 3D models, compared to 2D cell culture models, exhibit more *in vivo* tumour-like features including hypoxic regions, gradient distribution of chemical and biological factors and expression of pro-angiogenic and multidrug resistance proteins (4). These *in vivo* features are observed in 3D spheroids even though they are derived from a single immortal cell line. It is easy to envisage even more realistic 3D cell models, including cells embedded in active extracellular matrix (ECM) mimics (5) and incorporation of multiple cell types to form organoids (6). And yet 2D cell culture is still commonly employed (7) because of the ease of use and reproducibility of culturing cells on plastic or glass surfaces. Thus the challenge is how to produce large quantities of 3D cell cultures in an efficient and high-throughput manner such that statistically relevant data can be obtained (8).

For 3D cell culture to be used for both fundamental studies and drug screening (7) control over additional culture parameters is required. For example, it is known that volumes and shapes of spheroids influences the sensitivity to anti-cancer drugs (9, 10) and the ECM mimic that embed spheroids determines the rate of growth and drug responses (11, 12). There are a variety of methods for producing multicellular spheroids using either rotation, microfluidics, non-cell adherent surfaces, magnetic levitation or pealing confluent layers of cells off a surface (13-15), with the use of non-adherent surfaces being the popular approach (2). This method employs ultra-low attachment, round bottom well plates where gravity promotes cell aggregation to form a multicellular spheroid. To obtain the more *in vivo* realistic model of the spheroid embedded within an ECM mimic, requires the manual transfer of the spheroid into the appropriate tissue-like matrix (16), a process that is labour intensive and with a significant failure rate. An approach that overcomes the embedding challenge involves growing spheroids inside tissue-like hydrogels (17), such as Matrigel, collagen, alginate, synthetic peptides and polymers but at the cost of control over the spatial distribution and size of the spheroids. Thus, we urgently need a high-throughput method for producing embedded spheroids that is compatible with many cell types and ECM mimics and that provides control over spheroid size and shape so that statistically reliable data on drug responses can be obtained.

Recently, 3D bioprinting has emerged as a promising method to create reproducible but complex biological constructs by printing cell-laden hydrogel precursor or bioinks (18, 19). However, none of the existing 3D bioprinters have been designed specifically for producing 3D cell cultures and as such are not ideal for this application. Inkjet printing drop-on-demand technologies have high cell compatibility and has been used extensively to print geometrically simple hydrogel constructs (20, 21). Due to the droplet ejection mechanism, this technology shows very high cell viability but is limited to printing low viscosity bioinks (22) which effectively means bioinks with low cell concentration whereas high cell loadings are required for forming most 3D cell cultures. In contrast, the more popular microextrusion bioprinting process is used primarily in tissue engineering to print soft and hard tissues (23, 24). It is compatible with more viscous or semi-gel materials carrying a high density of cells. However, the imparted shear stress on the cells during printing affects cell viability (25).

Here, we establish a bespoke bioprinter and demonstrate the high-throughput bioprinting of embedded spheroids for various cell types. The printing process provided control over cell number, spheroid size and dimensions of the embedding cup without compromising cell viability. We performed extensive studies to confirm that the structural and biological characteristics of the 3D bioprinted spheroids were similar to manually prepared spheroids, including cell proliferation, apoptosis, cytoskeletal structure, hypoxia and stem-cell presentation. Further, we show that the bioprinting approach for embedded spheroid production enables high-throughput drug response analysis, measuring IC_50_ values for 12 conditions and 8 repeats in a 96-well plate. Our finding indicates that doxorubicin effectiveness depends on the size of the spheroid, whether it is embedded or not and how well the spheroid conforms to the embedding. In conclusion, our 3D bioprinter platform is a robust, reliable and high-throughput platform for embedded and non-embedded spheroids, overcoming many of the challenges of 3D cell culture models.

## Results

### 3D bioprinting of tumour spheroids inside a tissue-like matrix

The bespoke 3D bioprinter for the preparation of embedded 3D spheroids prints a high density of cells in a single droplet, with high viability directly into an extracellular matrix mimic (**Fig. 1A and Video S1**). The bespoke 3D bioprinter is comprised of a microvalve printhead, 2-axis robotics, a pressure regulation system, a printing stage for a 96-well microtiter plate and custom written software for biological assay design and printing. Here, alginate bioink and CaCl_2_ activator were used as the tissue-like matrix. Printing parameters, such as pressure, and bioink/activator concentrations were optimised to ensure that the biological properties of the ejected cells were not affected by fluidic shear stress or the ink materials. To examine whether printed cells remained viable, a trypan blue exclusion cell viability assay was performed to rapidly assess the viability of neuroblastoma (SK-N-BE(2)), non-small cell lung cancer (H460) and glioblastoma (U87vIII) cells (**Fig. 1B**). Cell viability of greater than 98% was observed for cells immediately prior to printing (pre-print) and after 3D bioprinting (post-print). The printing process also did not induce apoptotic cell death in the sample (**Fig. S1A**). To investigate the long-term biocompatibility of the tissue-like matrix, a live/dead assay was performed on the 3D encapsulated cells after 72 h incubation, which confirmed the biocompatibility of the chosen matrix, with viability of greater than 95% observed in all cases (**Fig. S1B-D**).

**Figure 1.**
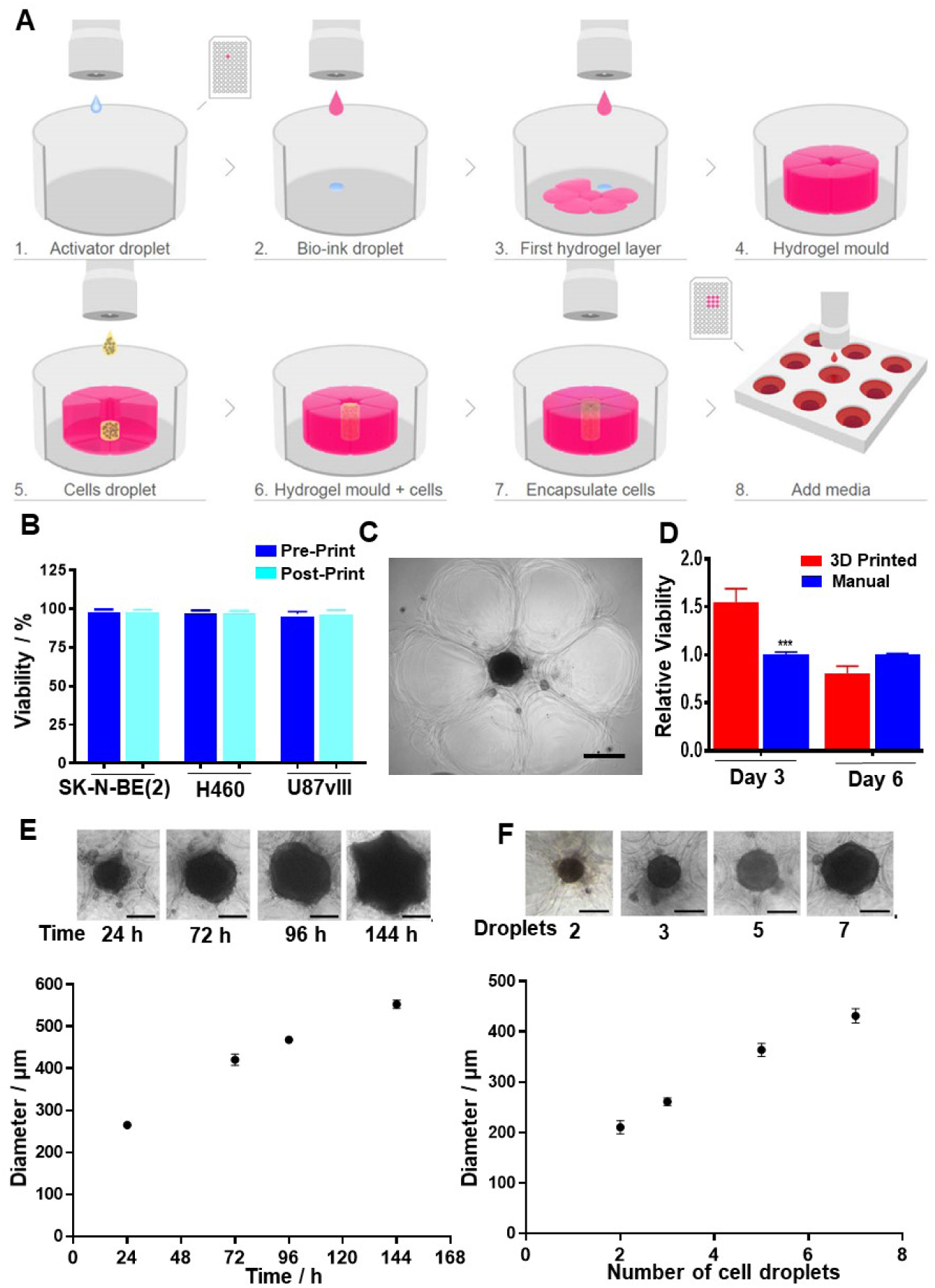
3D bioprinting of embedded spheroids. (**A**) A schematic of the bioprinting process. **Step 1-4**: A tissue-like matrix is bioprinted to form a cup (pink), **(Step 5-6)** into which the desired number of cells (yellow) are deposited. **Step 6-8**: The top-half of the cup is then bioprinted to complete the embedding and incubated at 37 °C and 5% CO_2_ until spheroids are formed (**B**) Cell viability of neuroblastoma (SK-N-BE(2)), non-small cell lung cancer (H460) and glioblastoma (U87vIII) cells in the bioink (Post-print) compared to cells before bioprinting (Pre-print) (*n* = 3) (**C**) Microscopy image of a bioprinted spheroid (middle black circle) surrounded by hydrogel matrix (outer flower-like structures). (**D**) Cell viability of bioprinted SK-N-BE(2) spheroids at day 3 and 6 compared to manually formed spheroids (*n* = 3, One way ANOVA; day 3 = *** P<0.0001, day 6 = non-significant (n.s.)). (**E**) The formation and growth of bioprinted SK-N-BE(2) spheroids over a period of 144 h with 23,750 cells (5 cell droplets) initially seeded. (**F**) Size of bioprinted SK-N-BE(2) spheroids after 3 days varied with the number of droplets containing cells that were seeded (**E-F**, *n* = 3 spheroids, scale bars = 200 µm). Results are means ± SEM.

To generate a single tumour spheroid embedded inside an alginate 3D hydrogel matrix structure, an alginate cup that had the structural stability to support gravity-based spheroid formation was printed (**Fig. 1A, steps 1-4 and Video S1**). Next, cell-laden ink at 250 million cells/mL was printed into the cup (**Fig. 1A, step 5-6**). The embedded spheroid was completed by printing the top layer of the hydrogel mould (**Fig. 1A, step 7)**. This process was then repeated; taking 80 minutes for printing an entire 96-well plate such that each well contained a single spheroid in a tissue-like matrix (**Fig 1A, step 8**). Upon incubation, a combination of gravitational forces and extracellular matrix (ECM) secreted by cells promoted cell migration, adhesion and proliferation and subsequent spheroid formation (2). Phase contrast microscope images of a printed spheroid of SK-N-BE(2) cells (**Fig. 1C**) showed that the cells formed a dense ball within the cup with the individual alginate droplets forming the cup (flower-like arrangement). The formation of the spheroid over a 72 h period are shown in **Video S2** and **Video S3** for the 3D bioprinted and manually prepared using a low-attachment, round-bottom well plate (26), respectively. The viability of SK-N-BE(2) cells within the spheroids were similar for printed and manually prepared spheroids after both 3 and 6 days (**Fig. 1D**) indicating neither the printing process nor the bioink were influencing the cells deleteriously. Similar observations were made for H460 and U87vIII cells (**Fig. S2A–B**).

The capability of the 3D bioprinter to accurately dispense discrete droplets with a high concentration of viable cells is critical for the formation of spheroids and unattainable using other printing technologies. Typically, five 19 nL droplets with approximately 4,750 cells per droplet were deposited inside the hydrogel mould. Monitoring the growth of the 3D bioprinted SK-N-BE(2) cells over 144 h (**Fig. 1E**) showed that the 3D bioprinted spheroids began to form after only 24 h. The rapid rate of spheroid formation was attributed to the high number of cells deposited inside a confined space. Similar observations were made for lung cancer cells (H460) and glioblastoma cells (U87vIII), respectively (**Fig. S2C-D**). All bioprinted spheroids increased in diameter linearly with time. Importantly, as the cup was completely filled by the growing spheroid, the spheroid began to conform to the shape of the cup which highlights the capability of the bioprinter to also produce spheroids with controlled shape by matching cup size, shape and cell volume. The size of the spheroids can be controlled via the cell density in each droplet or the number of droplets printed. It was demonstrated that increasing the number of SK-N-BE(2) cell droplets from two (∼9,500 cells) to seven (∼33,250 cells) led to an increase in spheroid diameter from 190 ± 13 µm to 420 ± 19 µm after 3 days of incubation (**Fig. 1F**). Thus, the 3D bioprinting platform enabled the rapid formation of embedded spheroids with high cell viability and control over spheroid size and shape.

### The in vivo tumour-like characteristics of the 3D bioprinted spheroids

We next addressed whether 3D printed matrix-embedded spheroids also possess important *in vivo* tumour-like characteristics found in manually prepared spheroids. To visualise the organisation of the 3D bioprinted SK-N-BE(2) spheroids, we immuno-stained for Ki67, a protein associated with cell proliferation (7), and stained the nucleus (**Fig. 2A**). Proliferating cells were consistently found on the periphery of the spheroid during the entire 6 days of investigation (**Fig. S3A**). Similarly, quantitative analysis of the light-sheet microscope images (**Fig. S3B**) located proliferating cells in both 3D bioprinted and manually prepared spheroids at the periphery (0.8-1 normalised distance, with 0 being the centre and 1 the edge of the spheroid). The presence of hypoxic cells in the 3D bioprinted spheroids was demonstrated using the well-established hypoxia marker, HIF1α. Dissociating and immuno-labelling the spheroids allowed analysis via fluorescence-activated cell sorting (FACS) which showed equivalent HIF1α positive cell populations of ∼5% in both the 3D bioprinted and manual spheroids (**Fig. 2B**), confirming that 3D bioprinted spheroids carry the important hypoxia characteristic of a 3D spheroid model (7). Manually prepared spheroids treated with CoCl_2_, an inducer of hypoxia, were used as positive controls for hypoxia (27).

**Figure 2.**
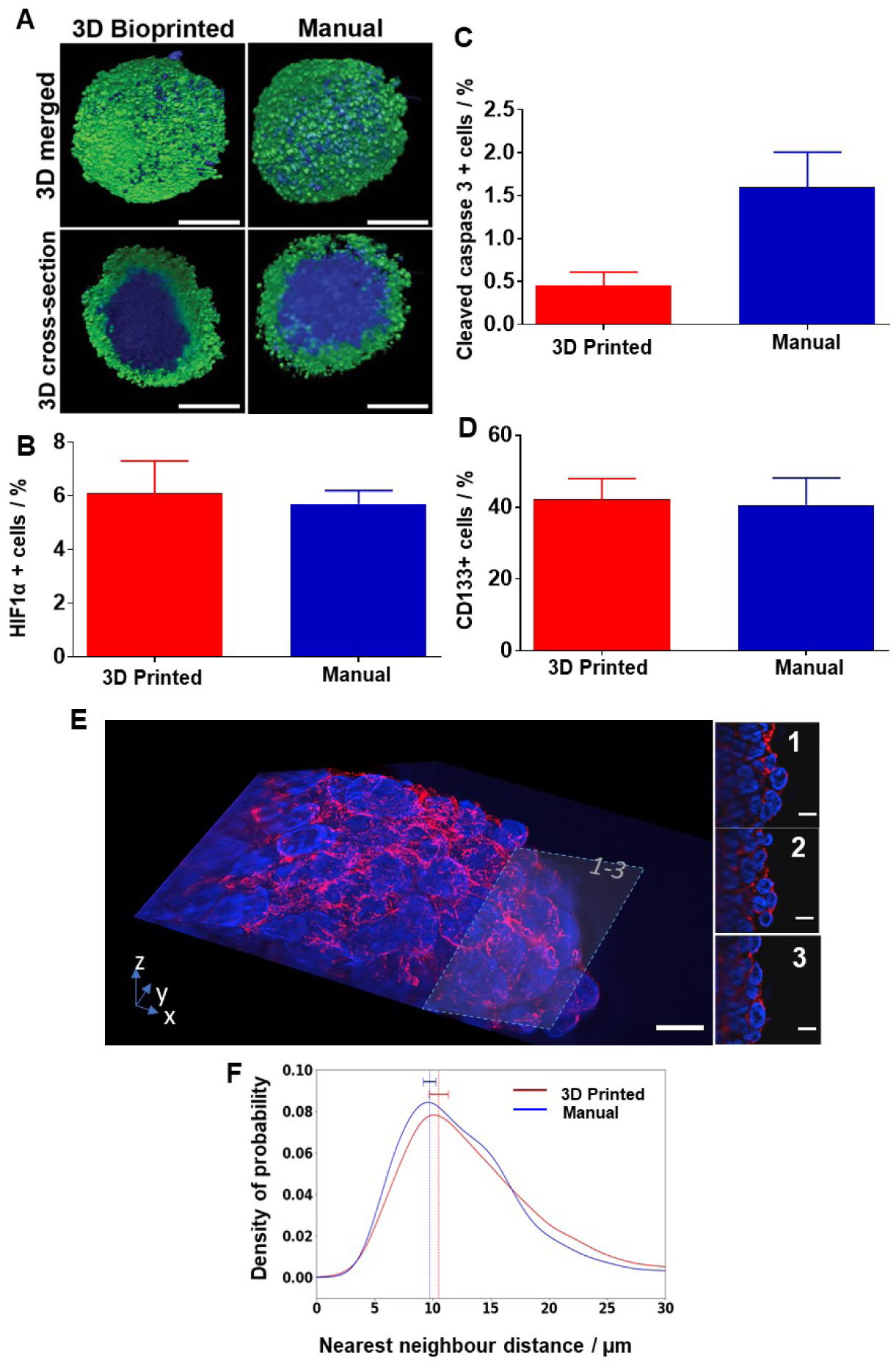
3D bioprinted spheroids have a similar organization as manually produced spheroids. (**A**) 3D rendered and 3D cross-sectioned (*via* optical sectioning) light-sheet microscopy images of the 3D bioprinted (left) and manual SK-N-BE(2) spheroids, labelled with α-Ki67 antibody (green), indicating cell proliferation and the DNA dye Hoechst 33342 (blue). Scale bars = 200 µm. (**B**) Percentage of HIF1α positive, (**C**) cleaved caspase-3 positive and (**D)** CD133 positive cells as determined by FACS. (In **B-D**, *n* = 3, Unpaired t-test; n.s.) (**E**) Lattice light-sheet images of a bioprinted spheroid, stained with phalloidin-568 (red) and SYTOX green (blue). The images were further sliced at 34.95 µm (**E1**), 27.45 µm (**E2**) and 19.95 µm (**E3**) from the top to show the high-resolution cellular arrangement of the spheroids. Scale bars = 10 µm. (**F**) Quantification of the cell-cell density was conducted by measuring the average distance between the nearest neighbouring nuclei in 3-dimension for both manually prepared and 3D bioprinted SK-N-BE(2) spheroids (*n* = 3). Results are means ± SEM

Next, we investigated apoptosis in 3D bioprinting spheroids. 3D bioprinted and manually prepared spheroids were dissociated and stained for the apoptotic marker cleaved caspase-3 prior to FACS analysis. Spheroids treated with doxorubicin (0.4 mM), were used as a positive control. The percentage of cleaved caspase-3 positive cells was 0.4% and 1.7% for the 3D bioprinted and manual spheroid, respectively, and not statistically significantly different (**Fig. 2C**). Apoptotic cells could also be seen in light-sheet microscopy images without a discernible difference in the number and location of cleaved caspase-3 positive cells between 3D bioprinted and manually assembled spheroids after 3 days and 6 days (**Fig. S4**).

We also assessed the percentage of cells with cancer stem like properties. These are key features that make *in vitro* 3D models more tumour-like than 2D models. Neuroblastoma cells express the cancer stem cell marker CD133, and CD133-positive neuroblastoma cells have the ability to form tumours(28). To determine whether CD133-positive cancer stem-like cells can be found in the 3D bioprinted spheroids, FACS analysis using anti-CD133 antibody were carried out. The results showed that nearly 40% of the cells of the 3D bioprinted and the manually prepared spheroids were CD133 positive (**Fig. 2D**), demonstrating the preservation of cancer-stemness in both types of spheroids.

To evaluate how the cells were arranged inside the 3D bioprinted spheroids, Hematoxylin and eosin (H&E) staining was performed on spheroid cross-sections. Micrographs of the sectioned spheroids showed that the cell arrangements and populations were very similar in both 3D bioprinted and manual spheroids at both day 3 and day 6 (**Fig. S5**). Lattice light-sheet microscopy of the spheroids stained with phalloidin for F-actin organisation (red) and SYTOX green for nuclei (blue) was used to explore the cell arrangement (**Fig. 2E)**. Cell-cell compactness was quantified by calculating the nearest neighbour distance for each nucleus in both 3D bioprinted and manual spheroids at day 3 with no significant difference was found (**Fig. 2F**). Thus, both qualitative 3D images and quantitative analysis confirmed that 3D bioprinting does not significantly alter the arrangement and compactness of cells within spheroids. In summary, we demonstrated that 3D bioprinted spheroids had the desired cellular organisation of cancer stem-like cells, apoptotic and hypoxic cells at the core and proliferating cells at the periphery.

Having shown that the compactness of 3D bioprinted and manual spheroids was similar, we investigated whether this was reflected in the drug penetration into spheroids. Drug penetration into 3D bioprinted spheroids (compared to manual spheroids) was performed using a previously reported method (29). To conduct a direct comparison between 3D bioprinted and manual spheroid, 3D bioprinted spheroid was recovered from the tissue-like matrix using the chelating solution. Both recovered 3D bioprinted and manually prepared spheroids were treated with 3.68 µM doxorubicin for 2 h. Doxorubicin diffusion into the spheroids was monitored with microscopy every 2 h for 10 h and quantified using a bespoke Python script. In brief, the image plane with maximum intensity for every time point was identified and used to calculate the penetration distance over time. For both types of spheroids, doxorubicin was only able to penetrate the periphery (**Fig. 3A**), which is attributed to the compact cellular arrangements, an observation that is commonly found in tumours. Over time, doxorubicin slowly penetrates and accumulates in the spheroids in both recovered 3D bioprinted and manual samples (**Fig. 3B**). These observations confirmed that the 3D bioprinted spheroids capture some of the biological behaviour of tumours, making them suitable for high-throughput 3D drug discovery experiments.

**Figure 3.**
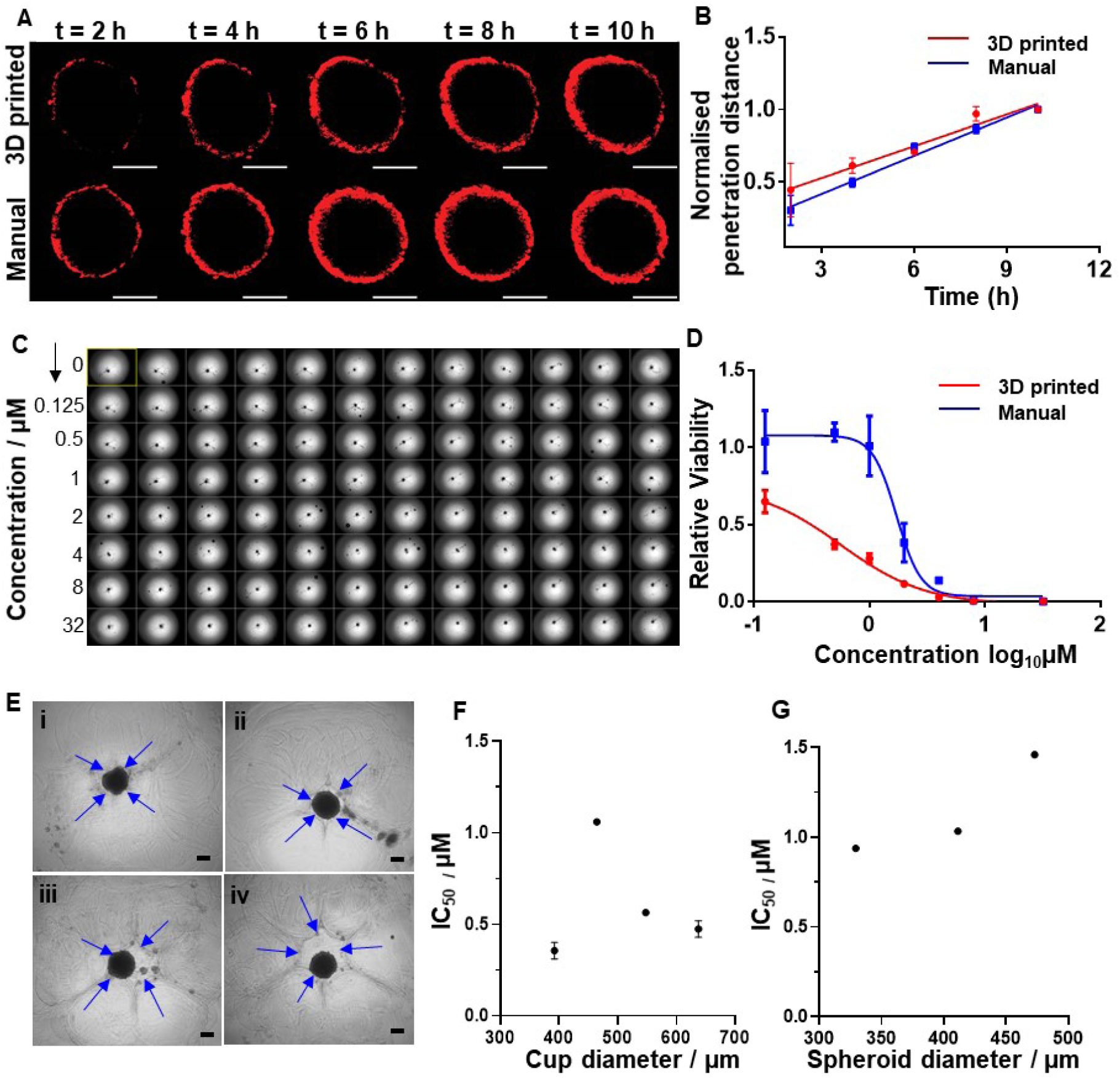
Bioprinted SK-N-BE(2) spheroids are efficient 3D models for high-throughput drug discovery (**A**) Confocal microscopy images at 2 h intervals over 10 h showing the penetration of doxorubicin (red) into the bioprinted and manually prepared spheroids. Scale bars = 200 µm. (**B**) Penetration depth profile of doxorubicin into bioprinted and manual spheroids over 10 h (*n* = 2, 3 spheroids group). (**C**) Operetta image of a full 96-well plate of 3D bioprinted spheroids, 12 replicates from each treated with 8 different doxorubicin concentrations. (**D**) Dose response curve of bioprinted spheroids, embedded in the hydrogel and non-embedded (*n* = 2, 6 spheroids per condition and per repeat). (**E**) Bioprinted spheroids encapsulated inside varying cup sizes. Spheroid diameter was kept similar at 400 ± 18 µm, while cup sizes was varied from (i) 376 ± 27 µm, (ii) 464 ± 29 µm (iii) 548 ± 27 µm and (iv) 637 ± 20 µm. Blue arrows indicate the cup wall. Scale bars = 200 µm). (**F**) Effect of the hydrogel cup diameter on IC_50_ of doxorubicin on bioprinted spheroids. (*n* = 2. 6 spheroids per group). (**G**) Effect of spheroid size on IC_50_ of doxorubicin on bioprinted spheroids with a constant cup size (*n* = 3 experimental repeats per condition). Results are means ± SEM.

### 3D bioprinted spheroids as a high-throughput tool for evaluating drug responses

The capability of 3D bioprinting of spheroids affords the opportunity of high-throughput drug screening. To show compatibility of the 3D bioprinted matrix embedded single spheroids for high-throughput screening, we exposed a 96-well plate of neuroblastoma spheroids, grown to an average diameter of 400 µm (72 h incubation post printing), to different concentrations of doxorubicin (0 - 32 μM) for a further 72 h (**Fig. 3C**). The drug response was measured directly in the microtiter plate using the CellTiter-Glo® assay, from which robust IC_50_ value was derived (**Fig. 3D**). The quality of the 3D bioprinted spheroids was tested with the statistically robust standardized mean difference (SSMD) analysis (30, 31). We conducted this analysis on all high-throughput screening experiments, with the results indicating that 95% of our printed spheroids, and hence the assays, were of good or greater quality, when compared to the acceptable criteria of the SSMD analysis (**Fig. S6**).

We next determined whether the embedding matrix itself has an impact on the drug response of spheroids, a parameter that has not been widely investigated. A direct comparison of the drug response was conducted between 3D bioprinted spheroids embedded in the hydrogel matrix and 3D bioprinted spheroids recovered from the hydrogel matrix. Interestingly, the presence of a matrix increased doxorubicin sensitivity of the spheroids (**Fig. 3D**) compared to free floating spheroids. This is possibly due to the matrix acting as a sink for the drug, thus accumulating the drugs from the media into the inner gel region. This in turn increases the availability of the drug to the spheroid thus increasing the drug efficacy.

As the previous experiment highlighted the importance of the hydrogel matrix on drug efficacy, we next looked at another variable, the size of the hydrogel cup in which the cells are deposited, and the spheroid grows (**Fig. 3E**). Four different cup sizes were bioprinted while keeping the spheroid diameter constant such that the space between the spheroid and the embedding hydrogel varied. Surprisingly, doxorubicin IC_50_ values were highly sensitive to cup size, indicating that matrix properties are an integral part of cancer biology (**Fig. 3F**).

Finally, by exploiting the capability of the 3D bioprinter to adjust spheroid size on demand (**Fig. 3G**), we quantified the effect of SK-N-BE(2) spheroid size on the drug response. Previous qualitative data suggest that MCF-7 spheroid size influences drug resistance(32). Three different spheroid diameters were generated, 329 ± 18 µm, 411 ± 32 µm and 473 ± 32 µm, by changing the number of cell droplets deposited into the cup (3, 5, and 7 droplets, respectively). We found an increase in doxorubicin IC_50_ values ranging from 1.06 to 1.48 µM as the spheroid sizes increase (**Fig. 3G**). Our data thus indicated that doxorubicin responses were inversely proportional to the surface area of the spheroids. The 3D bioprinting system presented herein is the ideal platform to create appropriate in vitro 3D cancer models and screen drug responses in a high-throughput manner.

## Discussion

Despite the importance of 3D cancer cell models and 3D extracellular matrix mimics has been well-established, yet their use in cancer research is not as widely adopted as it could be because of the difficulties involved in scaling up production of reproducible 3D cancer models embedded in a matrix. For the purpose of high-throughput and reliable production of 3D tumour spheroids, we developed a bespoke 3D bioprinter capable of printing droplets with cell concentrations up to 250 million cells/mL that maintains cell viability at greater than 98%. The combination of the chosen bioinks and non-contact, drop-on-demand, microvalve-based bioprinting technology used herein, allowed the printing of high cell density bioinks without exposing the cells to detrimental levels of shear stress during the droplet ejection process. The droplet-based bioprinting is easily scalable, in a manner analogous to inkjet printing, and facilitates simultaneous production of multiple embedded, multicellular, spheroid-containing 3D tissue culture models, making it suitable for rapid production of 3D *in vitro* samples. Using this technology, 3D multicellular spheroids encapsulated in a matrix were consistently printed into individual wells on a 96-well plate, demonstrating a novel method for high-throughput embedded 3D spheroid production. The bioprinted spheroids exhibited identical biological and architectural properties to manually prepared spheroids including overall viability, apoptosis, proliferation, cancer stemness and compactness was conducted to confirm that they can be used to replace widely used, labour-intensive manual spheroid culture methods.

Using the high-throughput bioprinting of spheroids we began a process of determining some of the important parameters in defining the drug response of spheroids; a thus far under explored aspect of 3D cell cultures. Our results demonstrate that the larger the spheroid the higher the IC_50_ for the model drug doxorubicin. Furthermore, the embedding process was shown to make the spheroid more sensitive to the drug, presumably because the matrix used herein accumulated the drug. Finally, we showed that the spheroid shape formed by the contact area between spheroid and ECM mimic impacted on doxorubicin effectiveness. Taken together, the capability of 3D bioprinted spheroids opens up many opportunities to create more relevant 3D cancer models in a high-throughput manner. In particular, primary cell mixtures and stem cells can be incorporated, which is of particular interest to precision medicine and regenerative medicine. The most powerful aspect of this 3D bioprinting platform is the ability for end users, such as cancer researchers, to intuitively design innovative assays, which will open up endless possibilities that can be expected to revolutionise biomedical research, including drug research and development.

## Materials and Methods

### Materials

Dulbecco’s Phosphate Buffer Saline (no calcium, no magnesium, DPBS, ThermoFisher Scientific), Dulbecco’s Modified Eagle Medium (DMEM, ThermoFisher Scientific), Roswell Park Memorial Institute 1640 (RPMI, ThermoFisher Scientific), Fetal Calf Serum (FCS, Bovogen), Penicillin-Streptomycin (Pen-Strep, 10,000 U/mL, ThermoFisher Scientific), trypsin-EDTA (ThermoFisher Scientific), ethanol (Univar), Triton-X (Sigma Aldrich), Bovine Serum Albumin (BSA, Sigma Aldrich) were used as received.

### Bespoke 3D bioprinter

3D cell culture models were bioprinted using a non-contact drop-on-demand 3D bioprinter (now manufactured by Inventia Life Science). The bioprinter incorporates a flyby or scanning printhead that prints directly onto the substrate while in motion, leading to high-throughput printing of 3D cell culture models. It comprises of a 2-axis motion control system, a droplet dispensing system and a pressure regulation system to control pressure in the fluid reservoirs. The bioprinter facilitates printing onto many types of multi-well plates. The 2-axis linear motion positioning system is capable of accurately positioning droplets on the substrate at a resolution of 20 μm along each axis. The droplet dispensing system consists of multiple independently addressable microvalves. Inks are primed into the nozzles from a bioprinting cartridge. The desired droplet volume can also be adjusted using the backpressure in the fluid reservoir and the microvalve opening time.

### Preparation of the bioink, activator, cell-carrier ink and chelating solutions

Three different ink solutions were used to produce the 3D spheroid assay. The bioink was prepared by mixing sodium alginate (FMC BioPolymer) in a 70/30 v/v% mixture of Milli-Q water and DPBS at 2 wt% for 16 h. The activator was prepared by mixing calcium chloride (Anhydrous, Sigma Aldrich) in Milli-Q water at 4 wt%. Lastly, the cell-carrier ink was prepared by mixing Ficoll 400 (Sigma Aldrich) in DPBS at 14 wt%. To make the chelating solution, sodium citrate (Sigma Aldrich) was dissolved in DPBS at 200 mM. All solutions were sterilised by filtration through a 0.22 µm syringe filter prior to use. The bioink was stored at 4 °C, while other solutions were stored at room temperature.

### Cell culture

SK-N-BE(2) (human neuroblastoma) and U87vIII (human glioblastoma) cells were maintained in 10% fetal calf serum (FCS)/ DMEM at 37 °C/5% CO_2_. Human non-small cell lung cancer H460 cells were cultured in 10% fetal calf serum (FCS)/RPMI at 37 °C/5% CO_2_. Cell lines are routinely screened and free of mycoplasma contamination.

### Cell viability and cell death assays

Trypan blue exclusion study was used to assess the viability of cells pre and post printing. SK-N-BE(2), H460 or U87vIII cells were harvested from 6-well plates using trypsin-EDTA and pelleted using centrifugation at 423 g. Cells were resuspended in bioink to a concentration of 2 x 10^8^ cells/mL. The printed samples were cells passed through the 3D bioprinter while non-printed referred to manually pipetted cells. Dead cells were identified using 0.4% trypan blue stain (Invitrogen). Cell counts were obtained by counting the number of live and dead cells in each sample. Percentage cell viability was determined by dividing the total number of live cells by the total number of cells (live + dead).

To investigate the potential impact of the printing on apoptotic cell death, SK-N-BE(2), H460 or U87vIII cells were resuspended in a bioink to a concentration of 2 x 10^8^ cells/mL. As controls, SK-N-BE(2), H460 or U87vIII cells were seeded into 6-well plates at 2 x 10^6^ cells/well and incubated at 37°C/5% CO_2_ for 24 h and 48 h. Cells pre- and post-printing were harvested as described above and stained with PE Annexin V (BD Biosciences), 7-aminoactinomycin D (7-AAD) (BD Biosciences), both PE Annexin V and 7-ADD or neither for 15 min at 37 °C in the dark. Apoptosis levels were measured by flow cytometry and analysed using FlowJo.

### 3D bioprinting of cancer spheroids

Bioink, activator and cell-carrier ink were brought up to room temperature prior to printing. 3D bioprinter surfaces were sterilised by 70% ethanol wiping. The cartridges, fluidics and microvalves were sterilised by 70% ethanol (10 mL) and sterile Milli-Q water (10 mL) flushing. The bioink (1 mL) and activator (1 mL) were then pipetted into their respective cartridges.

Bioprinting structure definition and printing execution were conducted using the proprietary custom-made software by Inventia Life Science. The bioprinting pressure set to 1.45 kPa and 0.55 kPa for the bioink and activator nozzle, respectively. Prior to initiating the printing process, spittooning was done on each microvalve to bring the inks into the nozzle and to ensure consistent ejection of liquid droplets. The hydrogel printing process was as follow: a drop of activator (18.5 nL) was initially printed at the desired location which was quickly followed by printing a drop of the complementing bioink (65.3 nL) at the same location to initiate the crosslinking process. This process was repeated until the desired alginate hydrogel structure was completed. Variation on the size of the 3D bioprinted cups was achieved by altering the microvalves opening time, in which larger opening time was used to achieve a smaller cup and vice versa.

Prior to printing the cells into the bioink, a cell-laden ink was prepared by dispersing 12.5 million cells of either SK-N-BE(2), H460, or U87vIII, in 50 μL of cell-carrier ink. The cell-laden ink was then pipetted into the printing cartridge, primed into the nozzle, pressurised to 0.59 kPa and spittooned. Five cell-laden ink droplets (19 nL each) were then bioprinted dropwise at 1 Hz into the hydrogel cup. Finally, the bioprinting process was completed by the printing of the top layer of the alginate hydrogel structure. At the completion of the printing process, 200 µL 10%FCS/DMEM (supplemented with Pen-Strep) was added manually in each individual well using a multi-channel pipette. The plate was then ready for incubation at 37 °C/5% CO_2_ for desired time depending on the experiment.

### Manual cancer spheroid formation

Manual spheroids were prepared using an ultra-low attachment (ULA) 96-well round-bottomed plates (Corning) using the previously described method (33). Briefly, the optimised cell number for each cell line (SK-N-BE(2), H460 or U87vIII) were pipetted into the ULA plate and cultured in their respective media (200 µL) for 3 or 6 days at 37 °C/5% CO_2_.

### 3D bioprinted spheroid recovery from the hydrogel matrix

Sterile chelating solution was used to recover the 3D bioprinted spheroids from the hydrogel matrix. The incubating media of the 3D bioprinted spheroids in the matrix, after the desired incubation time, was firstly removed using a multi-channel pipette. 100 µL of the chelating solution was added into each well and the sample was incubated for 2 min at room temperature to dissociate the matrix. After two minutes, the content of the well was carefully transferred into a microcentrifuge tube (Eppendorf) for further analysis.

### H&E staining

Both the 3D bioprinted and the manually prepared SK-N-BE(2) neuroblastoma spheroids were fixed in 4% paraformaldehyde (PFA, Electron Microscopy Sciences) diluted in 1% Triton (Sigma-Aldrich) in DPBS for 2 days at 4 °C. The fixed spheroids were first embedded in a small amount of alginate hydrogel to hold them in place in a Tissue-Tek cryomolds (VWR), which were subsequently filled with 500 µL of 4 w/v% agarose (AppliChem). Solidified blocks were transferred to 50% ethanol for 1 h and stored in 80% ethanol. Paraffin embedding, sectioning and H&E staining of spheroids were performed as previously described (26).

### CellTiter-Glo® 3D Viability assay

Two different variations of the CellTiter-Glo® 3D Viability assay (Promega) were used in this study. Assessment of the viability of manual spheroids and 3D bioprinted spheroids recovered from the hydrogel as per the manufacturer’s protocol in a 96-well solid white flat bottom plate (Corning). Briefly, spheroids were treated with 100 µL CellTiter-Glo® 3D solution in 96-well plates. Plates were incubated for 1 hour at room temperature on a Ratek microtiter PCR plate shaker at 30 rpm, prior to reading plates. Luminescence readings were obtained using the Perkin Elmer Victor3 plate reader. A standard built in protocol using appropriate filters were used to measure luminescence values. Average value of 3 luminescence readings were used as the final value. Spheroids treated with 0.4 mM doxorubicin (Teva Pharmaceuticals) for 1 h at room temperature were used as a positive control for cell death.

To consider the spheroid size variation between the 3D bioprinted and manual spheroids, the luminescence values were normalised to the volume of the spheroids. The volumes of the spheroids were calculated using the diameter of the spheroids obtained using a bright-field microscope and under the assumption that every spheroid was spherical. The spheroid volume-normalised luminescence values were further normalised to the manually generated spheroid.

In order to develop an HTS readout assay for our CellTiter-Glo® 3D protocol described above, we made slight modifications to the protocol. 25 µL of CellTiter-Glo® 3D solution was diluted in 175 µL of DPBS to obtain a total of 200 µL of final solution for each well of a 96-well plate. 3D bioprinted spheroids in 96-well optical-bottom black plates (Cell Carrier, Perkin Elmer) were tested in situ using the CellTiter-Glo® 3D assay. Luminescence readings were performed following the same protocol as described above.

### Drug penetration assay

To determine the ability of a fluorescent chemotherapy drug doxorubicin to penetrate spheroids, 3D bioprinted spheroids recovered from the hydrogel and the manually prepared spheroids were placed in an ultra-low attachment 96-well round bottom well plate (Corning). Spheroids were then incubated with 3.68 µM doxorubicin in 10%FCS/DMEM for 2 h at 37 °C. The spheroids were then imaged every 2 h for 10 h using ZEISS LSM 880 with Airyscan confocal microscope. Images were taken using 10x objective, Argon laser (488nm) and transmitted light (bright field). Z stacks were defined from the bottom to the middle of the spheroids. For every time point, an image stack on the z-axis was taken. The microscope incubator was set to 37 °C/5% CO_2_.

### Fluorescence activated cell sorting (FACS) analysis for quantitative cell analysis

We investigated the expression of different cellular biomarkers using FACS analysis. Spheroids were treated with trypsin-EDTA for 20 mins at room temperature to separate into single cells, followed by neutralisation of trypsin-EDTA using 10% FCS/DMEM. Cells were spun down at 423 g and washed once with DPBS. Cell fixing was done by firstly suspending cell pellets in 4% PFA for 20 mins at room temperature. Cells were then pelleted at 423 g and washed twice with DPBS.

Fixed cells were permeabilised with 0.1% Triton/DPBS for 10 mins at room temperature, washed twice with DPBS and blocked with 2%BSA in DPBS for 1 h at room temperature. α-cleaved caspase-3 (Cell Signalling Technology), α-HIF1α (abcam) or α-CD133 antibodies (abcam) diluted to 1/100 with 2% BSA/DPBS solution were incubated overnight with the cells. After removing excess primary antibody, the samples were stained with Alexa Fluor 488 labelled Goat anti-Rabbit IgG secondary antibody (ThermoFisher Scientific).

Positive staining was determined by flow cytometer BD FACSCanto using the 488 laser and analysed using FlowJo. Single cells were defined using Area Scaling Strategy (FCS-A vs FCS-H). Total Events (area based) were presented on the x-axis (FCS-A) and the events detected in the 488 nm laser channel were presented on the y-axis. Unstained cells used as negative control were used to set the baseline for the y-axis detected events and the percentages of positive cells above the baseline were shown in bar graphs as a direct comparison between manual and 3D printed cells.

### Immunofluorescence staining

Fixation and immunofluorescence protocols for light-sheet microscopy were adapted from a previously described protocol (34). To improve penetration of staining reagents to the core of the spheroids, simultaneous fixation and permeabilising was performed on 3D spheroids in 4% PFA/1% Triton-X/DPBS solution for 3 days at 4 °C. Spheroids were washed once with DPBS after removing the fixation media and stored in 0.1% Triton-X/DPBS at 4 °C. Fixed and permeabilised spheroids were blocked in 2% BSA in 0.1% Triton/DPBS for 3 days at 4 °C. To remove excess blocking solution, the samples were washed once with 0.1% Triton/DPBS.

To stain spheroids with α-Ki67/Hoechst 33342 or α-cleaved caspase3/Hoechst 33342, blocked spheroids were firstly incubated for 2 days at 4 °C with α-Ki67 (Abcam, stock concentration 0.031 mg/mL) or α-cleaved caspase3 (Cell Signalling Technology, stock concentration is not provided by the manufacturer) antibodies. The primary antibodies working solution was prepared by diluting the stock solution in 0.1% Triton/DPBS at 1/100. After incubation, excess primary antibodies were removed by washing the samples twice with 0.1% Triton/DPBS. Subsequently, the samples were incubated with 2 µg/mL Alexa Fluor 488 labelled Goat anti-Rabbit IgG secondary antibody (ThermoFisher Scientific, stock solution, 2mg/mL) prepared by mixing the stock solution in 0.1% Triton/DPBS at 1/100 and 0.2 mM Hoechst 33342 (ThermoFisher Scientific, stock concentration 20 mM) prepared by mixing the stock solution in 0.1% Triton/DPBS at 1/100, at 4 °C, overnight. Finally, the stained spheroids were washed twice with 0.1% Triton/DPBS.

To prepare phalloidin/Hoechst 33342 stained spheroids, blocked spheroids were firstly stained with phalloidin conjugated to Alexa Fluor® 568 (ThermoFisher Scientific) under vigorous shaking at 4 °C, overnight. Phalloidin working solution (0.31 µM) was prepared by mixing 5 µL of phalloidin conjugated to Alexa Fluor® 568 stock solution (stock concentration; 6.6 µM) in 100 µL of 0.1% Triton/DPBS. After incubation, the spheroids were washed twice with 0.1% Triton/DPBS to remove excess phalloidin. Nuclei were labelled by incubating phalloidin-labelled spheroids in Hoechst 33342 (0.2 mM) overnight at 4 °C on a shaker.

For Lattice light sheet imaging, the above method to prepare phalloidin-stained spheroids was followed to produce spheroids stained with phalloidin and. To stain the nuclei, phalloidin-stained spheroids were incubated in 0.167 µM SYTOX green (ThermoFisher Scientific) overnight at room temperature and protected from light. SYTOX green working solution was prepared by diluting SYTOX green (5 mM stock concentration) in DPBS at 1:30,000. After staining with SYTOX green, the spheroids were washed twice with DPBS and used immediately for lattice light sheet imaging.

### Light-sheet microscopy

The samples stained with appropriate markers were resuspended in 1% low melt agarose (AppliChem) and placed in ZEISS blue capillaries (size 4). ZEISS Lightsheet Z.1. (Carl Zeiss Microscopy GmbH, Jena, Germany) with 20x/1.0 (water, nd = 1.33) detection objective and two-sided 10x/0.2 illumination objectives equipped with two PCO EDGE 4.2 cameras was utilised to image two angles (0^0^ and 180^0^) of the spheroids from two sides (right and left). Laser lines 405 nm/50 mW, 488 nm/50 mW and 561 nm/50 mW were used with beam splitters/emission filters SBS LP 490/ BP 420-470, SBS LP 490/BP 505-545 and SBS LP 560/ LP 585 respectively. To fuse the right and the left sides of the images either manual dual side fusion or online dual side fusion was performed using Zen(black) (Carl Zeiss Microscopy GmbH) software package according to manufacturer’s instructions. To fuse the image stacks obtained from the two angles, software provided by the BioMedical Imaging Facility of University of New South Wales, Sydney, Australia was utilised.

### Lattice light-sheet microscopy

In order to investigate the compactness of cells within the spheroids we used lattice light sheet microscopy. Recovered 3D bioprinted spheroids, stained with phalloidin/SYTOX green prepared as described previously were carefully placed on top of a 0.01% poly-L-lysine (Sigma Aldrich) coated 5 mm round coverslip. Prepared samples were subsequently imaged using a lattice light-sheet microscope (Intelligent Imaging Innovations Inc.). The microscope generated 1 µm thick light-sheets with a square lattice configuration in dithering mode. Images were acquired with two ORCA-Flash 4.0 scientific complementary metal-oxide semiconductor (sCMOS) cameras (Hamamatsu). Each camera acquired a single colour channel sequentially to build up a respective 3-dimensional image. Imaging planes were exposed for 20 ms with 488 nm or 642 nm laser light. Spheroids were imaged on a piezo sample stage with 150 nm-step movement, thereby capturing a volume of ∼ 50 × 50 × 50 µm (512 × 512 × 333 pixels) within 20-25 s. The top edge of the spheroid was selected for assessing the imaging depth (∼ 50 µm). Acquired images were saved as TIFF files, with each plane of a volume corresponding to a cross-section of the imaged sample. The acquired images in the raw data appeared as a shear-transformed projection of the sample because the samples visualised in scan mode were scanned at a slanted angle to the coverslip of 31.5°. To compensate this coordinate distortion, images were de-skewed using inverse shear transform (Slidebook6, 3i). De-skewed volumes were then deconvolved with Slidebook using the constrained iterative deconvolution algorithm and an experimental point spread function generated with 200 nm fluorescent beads (ThermoFisher Scientific).

### Image analysis using Python

A python script employing the OpenCV Python binding library (35) and the python-bioformats library was used to analyse the image sets of the Hoechst fluorescent channel to obtain the coordinates of each fluorescent cells in a 3-dimensional space. For each *z*-layer planar image, the script equalised the images’ histogram, applied a Gaussian adaptive threshold using the “adaptiveThreshold” function and conducted a morphological closing transformation using the “morphologyEx” function to obtain a binary image of white blobs that represented the fluorescent cells. Into the binary image, a mask was applied and the “SimpleBlobDetector” function was used to find the bright cells surrounded by a black background. The coordinates at the centre of the blobs were recorded as the positions of the fluorescent cells’ centre of mass in the 2D *x*-*y* plane. As the process was repeated through all *z*-slices, repeated coordinates were removed. For the instance where repeated coordinates that represent one cell distributed in multiple *z*-slices, only the middle occurrence was kept to obtain the position of the cell centres of mass in 3D. The output of the analysis consisted of the recorded positions and sizes in µm, based on the OME-XML metadata conversion from pixel to µm, of the centres of all the detected blobs.

The detected coordinates of each fluorescent cells were processed using Python numerical and statistical analysis script, utilising Numpy, Pandas and Scipy libraries. Data points of the blobs were firstly filtered for outliers. We considered blobs that were too big or too small (compared to the median blob size) to be cells, and blobs that were detected outside of the region covered by the fluorescent cells (i.e. their distance from the centre of mass of the data points was too big or too small compared to the median distance) as the outliers. We used a median-deviation filtering technique to filter the outliers. For a statistical series *s = s*_*1*_,*…, s*_*n*_, which in this paper would be the series of the blob radii or the series of distances from the centre of mass, the deviation (*d*_m_) from the median value (median(*s*)), was defined as:

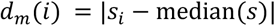

The modified z-score was calculated according to:

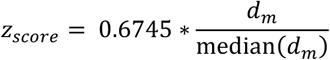

where median(*d*_*m*_) is the median of absolute deviation. To filter out the outliers, a maximum threshold value of 3.5 was applied as such only points of the series with z-score below 3.5 were retained (36).

To determine the spheroid compactness, the distance between nearest neighbour nuclei was calculated from the phalloidin/Hoechst light-sheet images. For each single point representing a Hoechst fluorescent cell nucleus, we computed the nearest neighbour from the remaining data points and recorded the corresponding distance. The resulting series contains the nearest neighbour distances for each cell (with 2 occurrences recorded for each cell, which doesn’t affect the actual distribution). The kernel density estimation of this series was then calculated using the gaussian_kde function from the Scipy.stats module. This function relies on a classic probability density functions (PDF) estimation based on the Gaussian kernels of bandwidth *n*^*-1/5*^ defined by Scott’s law for a 1-dimensional sample of size *n*.

The PDFs were plotted against the nearest-neighbour distance values to show the distribution of the nearest neighbour distances within each spheroid. The abscissa corresponding to the maximum of the PDF was recorded as a quantitative representation of the spheroid compactness, whereby a lower value represented a more compact spheroid. For both 3D bioprinted and manual spheroid sets, the global distribution functions were calculated as the arithmetic average of the individual spheroid’s PDF of the corresponding set. Therefore, we considered all spheroids as statistical entities and give them equal importance in the average. This was in contrast to a statistical analysis of the concatenated dataset of nearest neighbour distances that would skew the results towards the bigger spheroids. We also highlighted the average compactness and its variability in both 3D bioprinted and manual spheroid sets by calculating the average of the individual spheroids PDFs’ maxima and their standard deviation.

Locations of the proliferating cells in a spheroid were analysed from the Ki67/Hoechst stained light-sheet images. To compare the repartition of the Ki67 fluorescing cells between the inner region and periphery of the spheroids having variable geometries and dimensions, the data points were first re-centred around their centre of mass and projected to a sphere with a radius of 1 and centred at the origin (0, 0, 0). This was done by firstly subtracting the average (*x, y, z*) value from the data points. The resulting (*x, y, z*) coordinates were normalised against their respective maximum absolute value in both positive and negative directions along each axis according to:

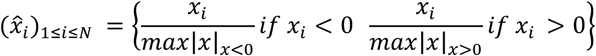

Where *I* is the index of each cell centre. The same formula was used for the *y* and *z*.

The distribution of the normalised points was then presented by calculating a kernel density estimation of their distance to the origin. The overall distribution for both 3D bioprinted and manual spheroid was plotted as an arithmetic average of each spheroid’s PDF within the corresponding 3D bioprinted or manual experimental set. This plot highlighted the repartition of the proliferating cells between inner region and periphery of each type of spheroid and its variability within each experimental set.

To quantify the doxorubicin penetration profile from the CZI microscope image set, comprising a set of bright-field and doxorubicin fluorescent images, we first selected the plane with the highest doxorubicin fluorescence intensity from the *z*-stack of the doxorubicin channel. Blob detection was then carried out on the corresponding bright field image to assign an approximated circular geometrical representation of the spheroid and to determine precisely the centre and the radius of the circle. After obtaining the geometrical characteristic of the spheroid slice, the biggest disc around the blob centre was extracted from the fluorescent image and divided into 100 concentric rings on which the average intensity was computed. The corresponding radial profile was then plotted at each time point to show the accumulation of drug inside the spheroid. The highest intensity reached at the final time point was recorded as a reference high value for the test. For each radial profile, we define the drug penetration distance as the distance between the maximum intensity (on the surface of the spheroid) and a minimum threshold defined as a significant fraction of the maximum. We choose 20% of the maximum as minimum threshold to obtain a strong enough intensity to be differentiated from the background. For comparison, penetration distance at each time point was normalised to the final value, and then averaged within both 3D bioprinted and manual spheroid sets. This produced the doxorubicin penetration distance plot.

### High-throughput 3D bioprinting of SK-N-BE(2) spheroids for high-throughput screening

SK-N-BE(2) spheroids for high-throughput screening (HTS) was bioprinted using the bespoke 3D bioprinter equipped with the proprietary fly-by printing logic, developed by Inventia Life Science, that enabled printing of twelve wells in a row simultaneously. For HTS of doxorubicin, a full 96-well plate (Cell-Carrier, Perkin Elmer) of SK-N-BE(2) spheroid inside hydrogel matrix were bioprinted according to the method detailed under the 3D bioprinting of cancer spheroids. Bioprinted spheroids were incubated in 10% FCS/DMEM (with Pen-Strep) at 37 °C/5% CO_2_ for 3 days before use. After incubation, the plate was imaged using Operetta (2x objective and bright field channel).

### High-throughput bioprinting of SK-N-BE(2) spheroids of different sizes

SK-N-BE(2) spheroids for this study were produced according to the bioprinting method described earlier with a slight modification. To generate the small spheroids, 3 droplets (19 nL/droplet) of cell-laden ink were printed, while 7 droplets (19 nL/droplet) of cell-laden ink were bioprinted to generate the large spheroids. The two different spheroid sizes were bioprinted into a 96-well plate (Cell-Carrier, Perkin Elmer), with 48 small spheroids bioprinted in row A-D and 48 large spheroids printed in row E-H. The plate was incubated for 3 days at 37 °C/5% CO_2_. After incubation, the plate was imaged using Operetta (2x objective and bright field channel).

### High-throughput bioprinting of spheroids in different hydrogel cup sizes

SK-N-BE(2) spheroids for this study were produced according to the bioprinting method described earlier. To vary the matrix enclosure size (or cup size) around the spheroid, the bioink microvalve opening time was varied, while keeping other printing parameters constant. The following opening time, 150, 135, 125 and 115 ck were used to generate a cup size around 380 nm, 450 nm, 550 nm and 630 nm, respectively. Five droplets of cell-laden ink (19 nL/droplet) were still printed into each cup. After bioprinting, 200 μL of 10% FCS/DMEM was added into each well and the plate was incubated for 3 days.

### HTS of Doxorubicin response of the bioprinted spheroids

Initially, Doxorubicin solutions at 0.125, 0.5, 1, 2, 4, 8 or 32 μM were prepared by mixing doxorubicin (Teva Pharmaceutical) in 10% FCS/DMEM. The solution was prepared fresh prior to each experiment. After incubating the bioprinted spheroids for 3 days, the incubating media was replaced with 200 μL doxorubicin solution. For every doxorubicin concentration, at least 3 wells were prepared. The plate was then incubated for 3 days.

For HTS of doxorubicin, a full 96-well plate of bioprinted spheroids were used to determine the doxorubicin IC_50_. After 3 days of incubation after bioprinting, the incubating media was replaced with 200 μL doxorubicin in 10% FCS/DMEM at various concentration, in the following arrangement: 0 μM in row A, 0.125 μM in row B, 0.5 μM in row C, 1 μM in row D, 2 μM in row E, 4 μM in row F, 8 μM in row G, 32 μM in row H. The plate was then incubated for 3 days. The effect of doxorubicin was then quantitatively assessed using CellTiter-Glo® endpoint analysis as described earlier. Dose response curve for each experiment was generated in GraphPad Prism software followed by calculation of IC_50_.

### Study of the effect of the hydrogel matrix on SK-N-BE(2) spheroid response to doxorubicin

Bioprinted SK-N-BE(2) spheroids were prepared as described earlier in a 96-well plate (Cell-Carrier, Perkin Elmer). After incubating the plate for 3 days, non-embedded bioprinted SK-N-BE(2) spheroid samples were prepared by recovering the spheroids from the matrix by following the method described earlier. The remaining embedded bioprinted spheroids in the well plate were exposed to 200 μL of doxorubicin in 10% FCS/DMEM at various concentration (0, 0.125, 0.5, 1, 2, 4, 8 or 32 μM). In a solid round bottom, white-walled 96-well plate (Costar), 200 μL of doxorubicin in 10% FCS/DMEM at various concentration (0, 0.125, 0.5, 1, 2, 4, 8 or 32 μM) was added. Into each well, one recovered 3D bioprinted spheroid was carefully transferred using a pipette. Both plates were incubated for 3 days at 37 °C/5% CO_2_. Quantification was then conducted using CellTiter-Glo® 3D endpoint readout using the method and analysis method as described earlier.

### Statistical analysis

Statistical analyses were performed using the GraphPad Prism v6 software (GraphPad Software). Unpaired, two-tailed Student’s t-tests were used to determine statistical differences between a control and an experimental group. For comparison of multiple samples, one-way ANOVA with a post hoc Bonferroni test was used. P-values of <0.05 were deemed statistically significant. Results were expressed as means of the number of independent experiments performed (at least 2) ± standard error of mean (SEM).

Prior to conducting strictly standardised mean difference (SSMD) analysis, visual inspection of the HTS data was conducted and outliers were omitted, whereby outliers were described as samples with either no spheroid present or more than one spheroid present. SSMD analysis was conducted according to the method described previously, according to:

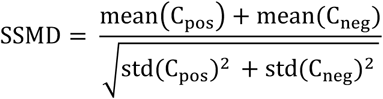

where, mean(C_pos_) and mean(C_neg_) were the mean of the positive and negative controls respectively and std(C_pos_) and std(C_neg_) were the standard deviation of the positive and negative controls respectively. For Exp 1 – 4, SSMD analysis was conducted on a minimum of 10 out of 12 samples (min. *n* = 10), while 6 out 6 samples (*n* = 6) were used for Exp 5 – 9.

## Data availability

Data available upon request to the corresponding authors.

## Author Contributions

R.H.U, L.A. and C.M.F. designed, performed and analysed experiments. L.A., A.P.O.M and T.A. established and performed the image analysis. J.R., A.P.O.M and K.J.O.M. designed and built the 3D bioprinter. J.B. performed and analysed the lattice light-sheet experiments. K.G. designed the lattice light-sheet experiments, contributed to the data analysis and interpretation. J.R., M.K. and J.J.G. designed the project strategy and experiments. All authors contributed to the writing of the manuscript.

## Acknowledgments

The authors would like to thank the Biomedical Imaging Facility (BMIF) within the Mark Wainwright Analytical Centre at UNSW for use of the facility and their technical assistance, in particular from Dr. Elvis Pandzic, Dr. Michael Carnell and Dr. Sandra Fok. The authors acknowledge the Children’s Cancer Institute, which is affiliated with the Sydney Children’s Hospital and School of Women’s & Children’s Health, UNSW Sydney. The authors acknowledge the following funding sources: Australian Research Council (ARC) Linkage Grant (LP130101035 to J.J.G., M.K., J.R.); ARC Laureate Fellowship (FL150100060 to J.J.G); National Health and Medical Research Council (NHMRC) Program Grant (APP1091261 to M.K, J.J.G); NHMRC Principal Research Fellowship (APP1119152 to M.K) and an NHMRC Senior Research Fellowship (1059278 to K.G.), support from ARC Centre of Excellence in Convergent Bio-Nano Science and Technology (CE140100036 to J.J.G and M.K) and ARC Centre of Excellence in Advanced Molecular Imaging (CE140100011 to K.G.).

## Notes

### Competing Interest Statement

A.P.O.M, T.A., K.J.O.M and J.R. are employees and/or consultants of Inventia Life Science Pty Ltd. Inventia Life Science Pty Ltd has an interest in commercializing the technology.

