## Supplementary Figures for "Precise, high-throughput production of multicellular spheroids with a bespoke 3D bioprinter"

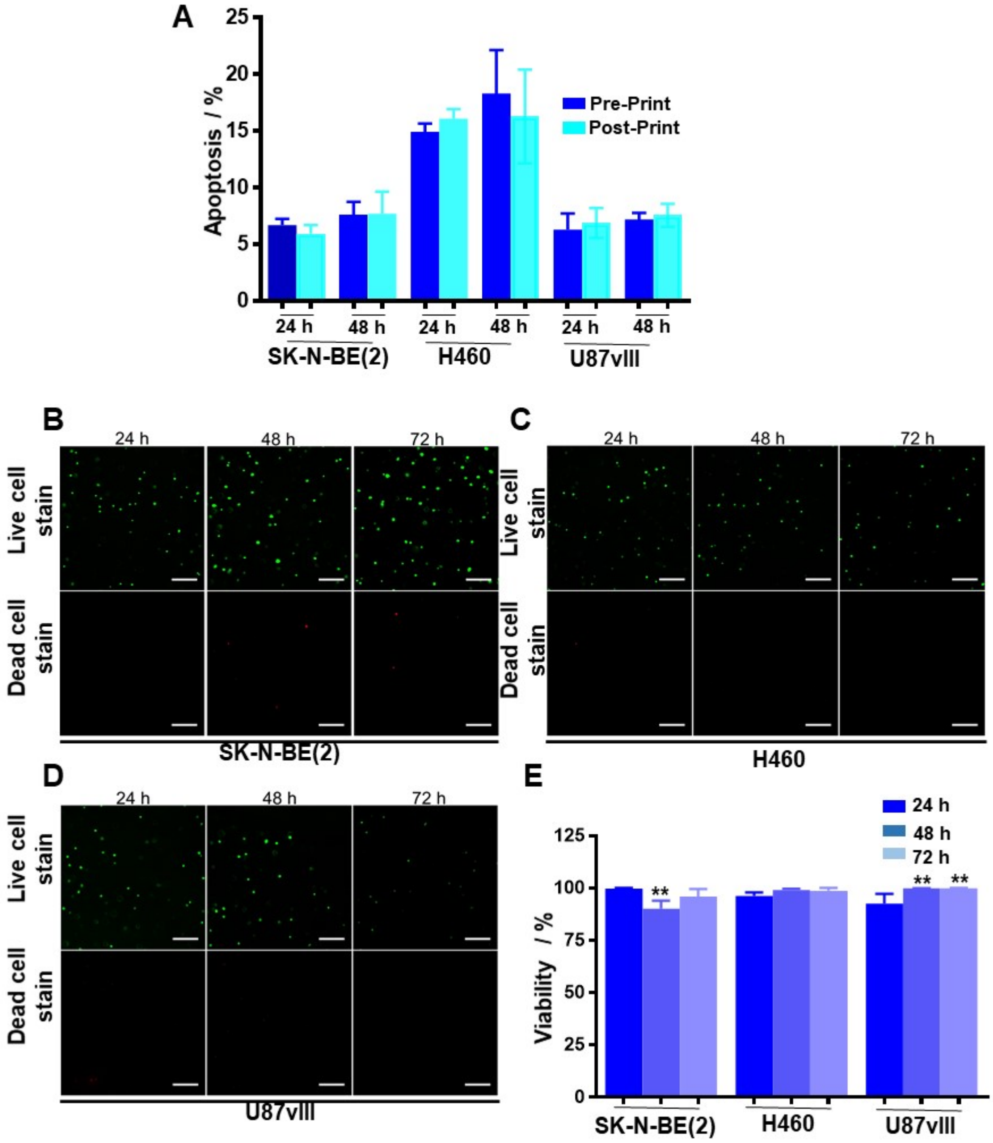


**Figure S1.** Cell viability and apoptosis studies on neuroblastoma (SK-N-BE(2)), non-small cell lung cancer (H460) and glioblastoma (U87vIII) cells (**A**) Annexin V apoptosis assay showing no visible effect of bioprinting on the cells when compared to the non-printed cells. 3D bioprinted neuroblastoma (SK-N-BE(2)), non-small cell lung cancer (H460) and glioblastoma (U87vIII) 3D cell encapsulation assay. Cells were dispersed in the bioink at 2 x $10^8$ cells/mL and bioprinted, together with the activator, to embed single cells inside the 3D tissue-like matrix. ($n$ = 3, One-way ANOVA; n.s.) (**B-D**) Live/dead assay staining on the samples after 24, 48, and 72 h incubation, using calcein AM (green) that stained living cells and ethidium homodimer (red), which stains dead cells. Scale bar = 250 µm. (**E**) Counted number of cells from the microscopy images showing that viability of > 95% was observed in all samples and across three different cell lines. ($n$ = 3, One-way ANOVA; SK-N-BE(2): **P < 0.05, H460: n.s., U87vIII: **P < 0.05). Results are means ± SEM.

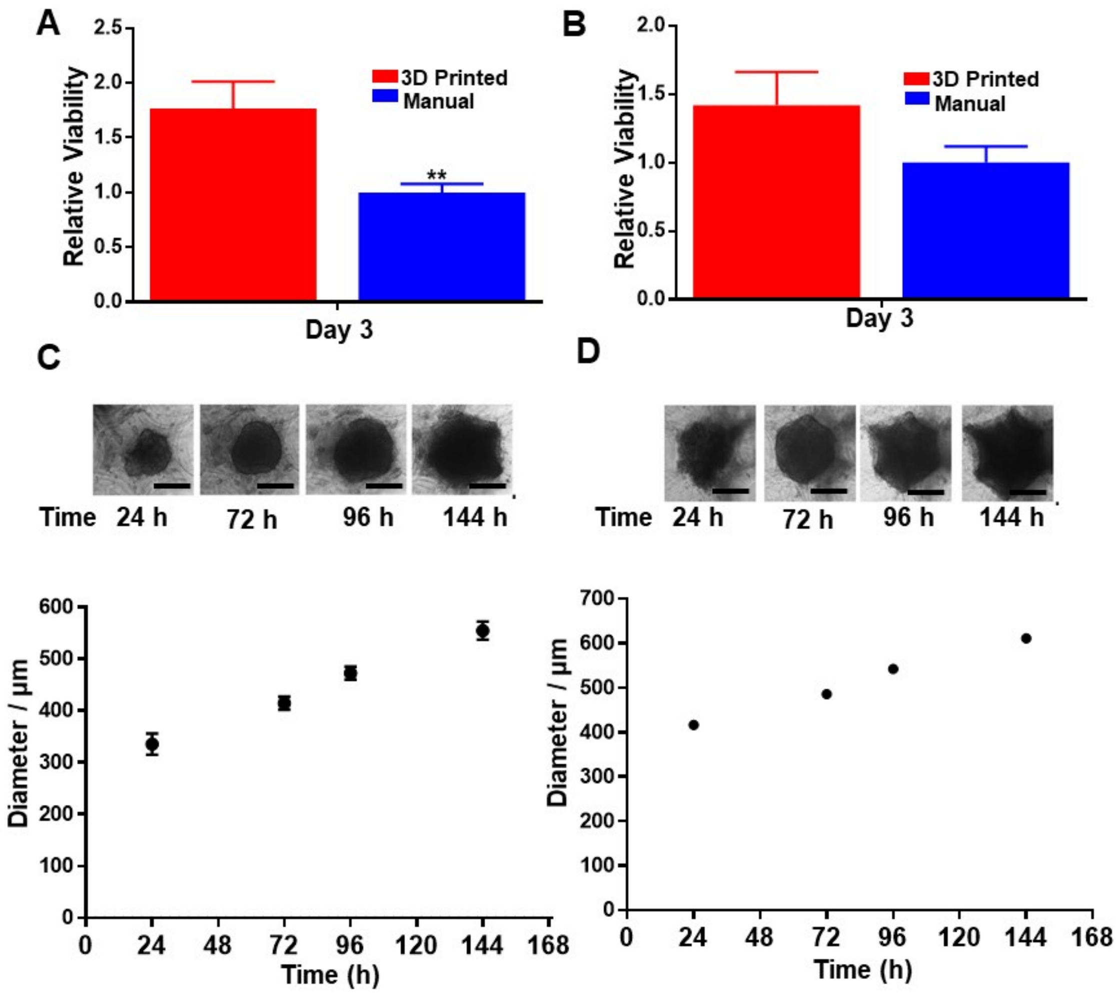


**Figure S2**. Spheroid viability and growth studies on non-small cell lung cancer (H460) and glioblastoma (U87vIII) cells (**A**) 3D bioprinted H460 showed significantly higher viability over manually prepared spheroids at day 3 ($n$ = 3, Paired t-test; **P < 0.05) (**B**) U87vIII spheroids and manually formed spheroids had similar viability at day 3 ($n$ = 3, Unpaired t-test; n.s.)(**C**) The formation and growth of H460 and (**D**) U87vIII spheroids over a period of 144 h from an estimated 23,750 cells printed for H460 and approximately 45,000 cells for U87vIII. In all cases, spheroids formed after 24 h and conformed to the shape of the matrix cup after 144 h. ($n$ = 6 experimental replicates, scale bars = 200 µm). Results are means ± SEM.

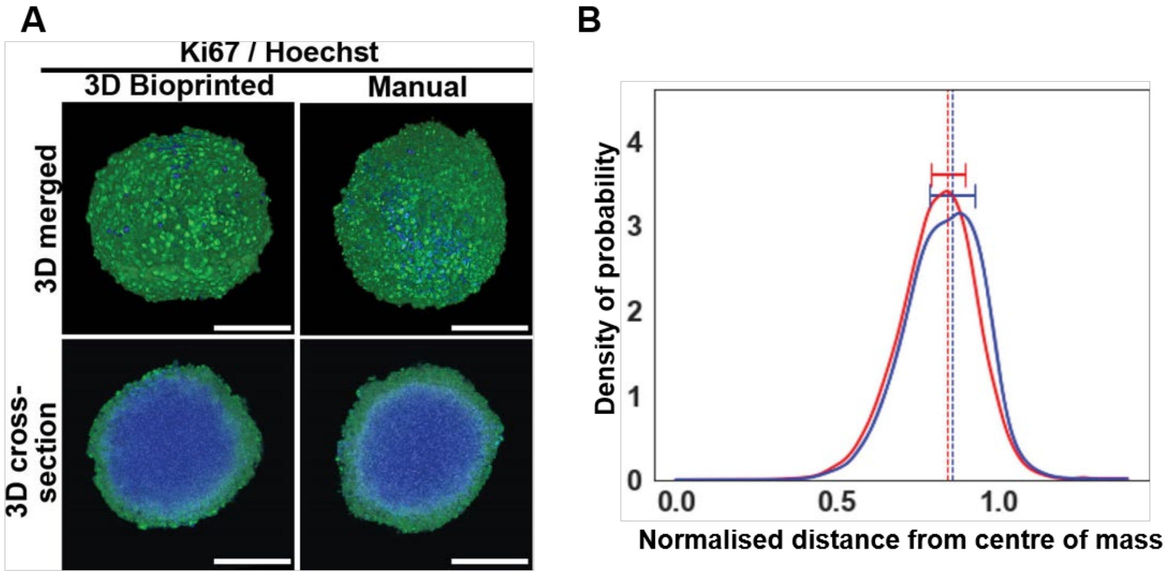


**Figure S3.** Ki-67 immunofluorescence analysis of the 3D bioprinted spheroids after 6 days of incubation. (**A**) Cells within the spheroids were labelled with α-Ki67 antibody (green), indicating cell proliferation and the DNA dye Hoechst 33342 (blue). The 3D cross-section images showed that the location of proliferating cells was consistent with the observed results on day 3 spheroids. (*n* = 2, scale bars = 300 μm) (**B**) Averaged, normalized distance distribution of the proliferating cells in both 3D bioprinted (red) and manual (blue) spheroid, with an x-axis value of 0 indicating the centre of the spheroid and 1 indicating the periphery of the spheroid. The dashed lines indicate the distance with maximum probability, with the standard deviation presented for both spheroid sets. Results are means ± SEM.

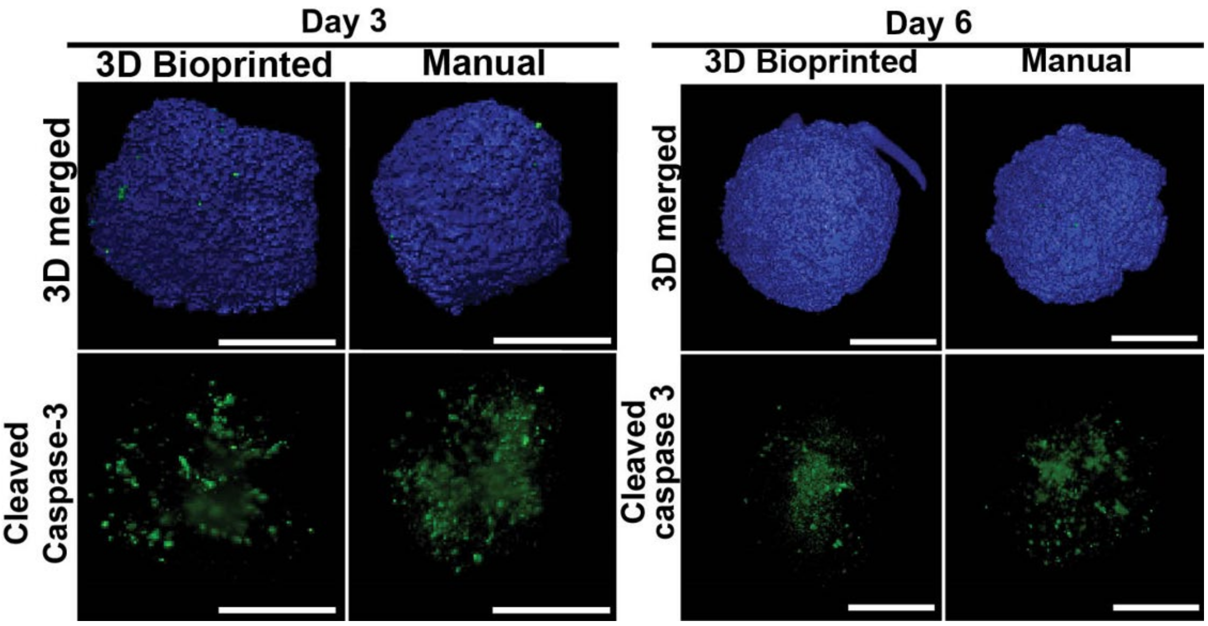


**Figure S4.** 3D rendered images of the 3D bioprinted and manual SK-N-BE(2) spheroids, labelled with α-cleaved caspase-3 antibody (green) and Hoechst 33342 (blue) and the corresponding 3D cleaved caspase-3 channel, after 3 and 6 days of incubation. The cleaved caspsase-3 channel images showing the location of apoptotic cells within the spheroids. Scale bars = 300 μm.

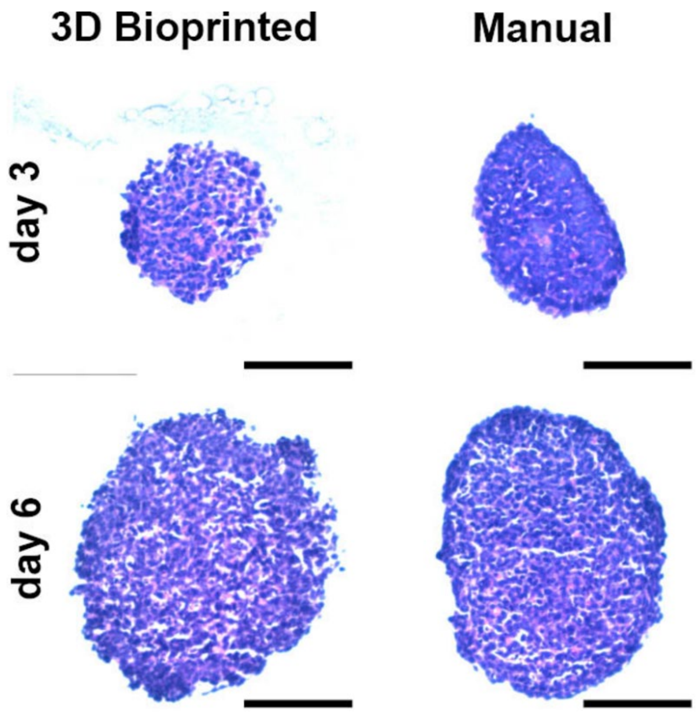


**Figure S5.** Micrographs of the H&E slices of the 3D bioprinted and manual SK-N-BE(2) spheroids at day 3 and day 6. Scale bars = 300 µm.

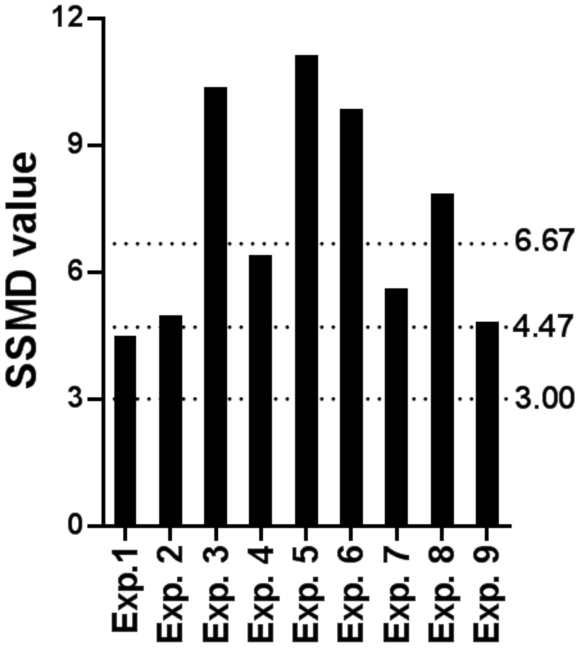


| SSMD | QC Interpretation |
|---|---|
| SSMD ≥ 6.67 | Excellent (E) |
| 6.67 > SSMD ≥ 4.7 | Good |
| 4.7 > SSMD ≥ 3 | Acceptable (A) |
| SSMD < 3 | Poor (P) |

**Figure S6.** Calculated SSMD values for each assay conducted for all HTS experiments, with the dotted line outlining the threshold value for the quality control (QC) interpretation. The corresponding table summarises the theoretical values of the quality control QC tests. For the SSMD value calculation, ($\underline{n}$ = *min. 6*).

**Video S1.** Video representation of the 3D bioprinting process of embedded 3D spheroids in a 96-well plate, depicting the bioprinting of the hydrogel cup structure and the deposition of high cell concentration droplets inside the cup.

**Video S2**. Time lapse video over 72 h representing the spheroid formation process of SK-N-BE(2) cells bioprinted inside the hydrogel cup.

**Video S3**. Time lapse video over 72 h representing the spheroid formation process of SK-N-BE(2) cells manually prepared using a low-attachment, round-bottom well plate.
